## Supplemental Table 1 for "A family of viral satellites manipulates invading virus gene expression and affects cholera toxin mobilization"

**Supplementary Table S1: Strains used in this study**

| Strain | Description* | Source |
| --- | --- | --- |
| KDS6 | *V. cholerae* E7946 O1, El Tor biotype | Lab collection |
| KDS36 | *V. cholerae* E7946 containing PLE1 | (1) |
| KDS37 | *V. cholerae* E7946 containing PLE2 | (1) |
| KDS38 | *V. cholerae* E7946 containing PLE3 | (1) |
| KDS39 | *V. cholerae* E7946 containing PLE4 | (1) |
| KDS40 | *V. cholerae* E7946 containing PLE5 | (1) |
| KDS281 | *V. cholerae* E7946 *∆ctxAB*::KanR | This study |
| KDS282 | *V. cholerae* E7946 *∆ctxAB*::KanR containing PLE1 | This study |
| KDS283 | *V. cholerae* E7946 *∆ctxAB*::KanR containing PLE2 | This study |
| KDS284 | *V. cholerae* E7946 ∆CTXφ, ∆lacz::P_tac_-*toxT,* SpecR | This study |
| ICP1 | ICP1_2006_E ΔCRISPR ΔCas2-3 | (2) |

* KanR = Kanamycin resistance cassette, SpecR = Spectinomycin resistance cassette.
